# iLAM: imaging Locomotor Activity Monitor for circadian phenotyping of large-bodied flying insects

**DOI:** 10.1101/2023.11.20.567947

**Authors:** Jacob N. Dayton, Avalon C.S. Owens

## Abstract

**Practical Tools:**

1. Historically, most insect chronoecological research has used direct observations, cameras, or infrared beam-based monitors to quantify movement across timed intervals. Although some alternative DIY systems are cheaper than the current standard locomotor activity monitor, these options remain complicated to build and/or computationally intensive.
2. We developed the **i**maging **L**ocomotor **A**ctivity **M**onitor (**iLAM**), an affordable (∼ $75 USD/unit) system for activity quantification. The iLAM utilizes a Raspberry Pi Zero W computer and night-vision camera inside a flight cage to photograph a population of insects at user-defined intervals. Open-source, modular R-scripts process the images and output a file containing the number, size, coordinate location, and timing for all movements (blobs) identified between consecutive images. Output can be analyzed directly or converted into the standard TriKinetics DAM format.
3. We demonstrated the flexibility and power of the iLAM system by comparing diel and circadian activity of different insect species (fireflies: *Photinus marginellus, P. greeni, P. obscurellus*), ecotypes (moths: *Ostrinia nubilalis*), and sexes (moths: *O. nubilalis*). Data captured by only six iLAMs ($450) identified that peak activity of *O. nubilalis* females (AZT: 19.2 hr) occurs significantly earlier than males (22.0 hr). Additionally, male moths from a univoltine population exhibited a significantly shorter endogenous period length (AZT: 21.3 hr) than males from a bivoltine genetic background (22.6 hr).
4. The iLAM will serve as a valuable tool for future researchers seeking to measure locomotor activity across diverse species, sexes, and populations in constant and changing environments.

## 1. Introduction

Almost all organisms on earth perform various tasks at different times of day to take advantage of variation in resource availability. The evolutionary and physiological drivers of these diel activity rhythms, as well as their behavioral consequences, have been a focus of chronobiological research for close to three centuries (Pittendrigh, 1993). While some activity patterns are directly cued by external signals such as light or temperature, others are endogenously driven by a highly conserved, temperature-compensated circadian clock mechanism (Sauders et al., 2002; Brady, Saviane, Cappellozza, & Sandrelli, 2021). In insects and other animals, the circadian clock sits atop of a hierarchy of time-variant processes as diverse and fundamental as vision, navigation, learning, metabolism, and development (Tomioka, Uryu, Kamae, Umezaki, & Yoshii, 2012). The potential for anthropogenic stressors to alter diel activity rhythms to the detriment of affected taxa has become a focus of recent research in chronoecology (Gaynor, Hojnowski, Carter, & Brashares, 2018; Levy, Dayan, Porter, & Kronfeld-Schor, 2019; Gilbert et al., 2022).

As chronobiological research writ large requires sampling consistently over extended periods (e.g. counting firefly flashes “every half-hour or hour for 28 consecutive hours”; Buck, 1937), much of which occurs outside of a typical workday, early advances in automated sampling were eagerly adopted (Konopka & Benzer, 1971). In the past two decades, Drosophila/Locomotor Activity Monitors (DAM/LAMs, hereafter referred to jointly as DAMs; TriKinetics, Waltham, MA) have become the primary method for recording the activity patterns of fruit flies and other small insects (Blanchardon et al., 2001). DAMs measure the timing and number of infrared beam interruptions made by individual flies moving across small glass tubes. DAMs have directly led to multiple breakthroughs in our understanding of the genetic and neuronal architecture of the circadian clock network in *Drosophila melanogaster* (Patke, Young, & Axelrod, 2020).

In addition to their high cost ($525-$3450), DAMs have several limitations that prevent their use with non-traditional model organisms (e.g., *Danaus plexippus*, Zhang, Iiams, Menet, Hardin, & Merlin, 2022). First, the small tubes (10-25 mm diameter) cannot easily accommodate large-bodied insects; they limit movement and preclude ecologically relevant behaviors such as flight, orientation, mating, and foraging. Second, recording duration in DAMs is limited because these tubes lack space for water or nectar supplementation (but see Wang, Yang, & Chen, 2021). Third, they reduce the complexity of ecologically relevant behaviors into binary beam interruptions which removes information otherwise captured by live observations or recorded images.

In recent years, automatic processing of video or image time series has emerged as an attractive alternative to beam-crossing activity monitors. However, the majority of these approaches are intended not for activity monitoring *per se* but for automatically tracking individuals, recording movement paths (Panadeiro, Rodriguez, Henry, Wlodkowic, & Andersson, 2021), and classifying behaviors (e.g., Bohnslav et al., 2021). These programs can achieve impressive results, but many of their higher-level capabilities – although reducible to gross activity over a 24-hour period – exceed the demands of chronobiologists and behavioral ecologists. Furthermore, the data storage and supercomputing infrastructure needed to process days of video pose barriers to many researchers and limit the scope of experiments (i.e., number of samples and recording duration). A less technically demanding and computationally intensive system explicitly designed for automated activity monitoring would increase the accessibility of this technique and help bring chronoecological research into the modern era.

Our **i**maging **L**ocomotor **A**ctivity **M**onitor (iLAM) employs an infrared Raspberry Pi camera mounted on the bottom of a DIY flight cage. The camera periodically takes images of large-bodied insects flying or crawling overhead. Images are saved onto an SD card where they can be downloaded and automatically analyzed with scripts written in the programming language R. The scripts subtract consecutive pairs of images, identify differences between them, and then record where these differences – i.e., movements – occurred, how large they were, and when they took place.

In this paper, we describe how to build and operate an iLAM and demonstrate its use in three case studies: one on the temporal niches of *Photinus* fireflies and the other two on sex-specific and ecotype differences in activity timing of the European corn borer (*Ostrinia nubilalis*). We conclude by comparing our iLAM system with other activity monitors currently in common use.

## 2. Materials and Methods

### 2.1 iLAM Design and Construction

Detailed step-by-step instructions for iLAM construction, image processing, activity analysis, and troubleshooting are provided online as Supporting Information (https://[xxx].github.io/ilam/). Briefly, the iLAM consists of three main components: (1) flight cages to contain insects, (2) single-board computers connected to night-vision cameras for imaging, and (3) customizable scripts to control imaging frequency, file export, and post-process image segmentation (Figure 1).

**Figure 1.**
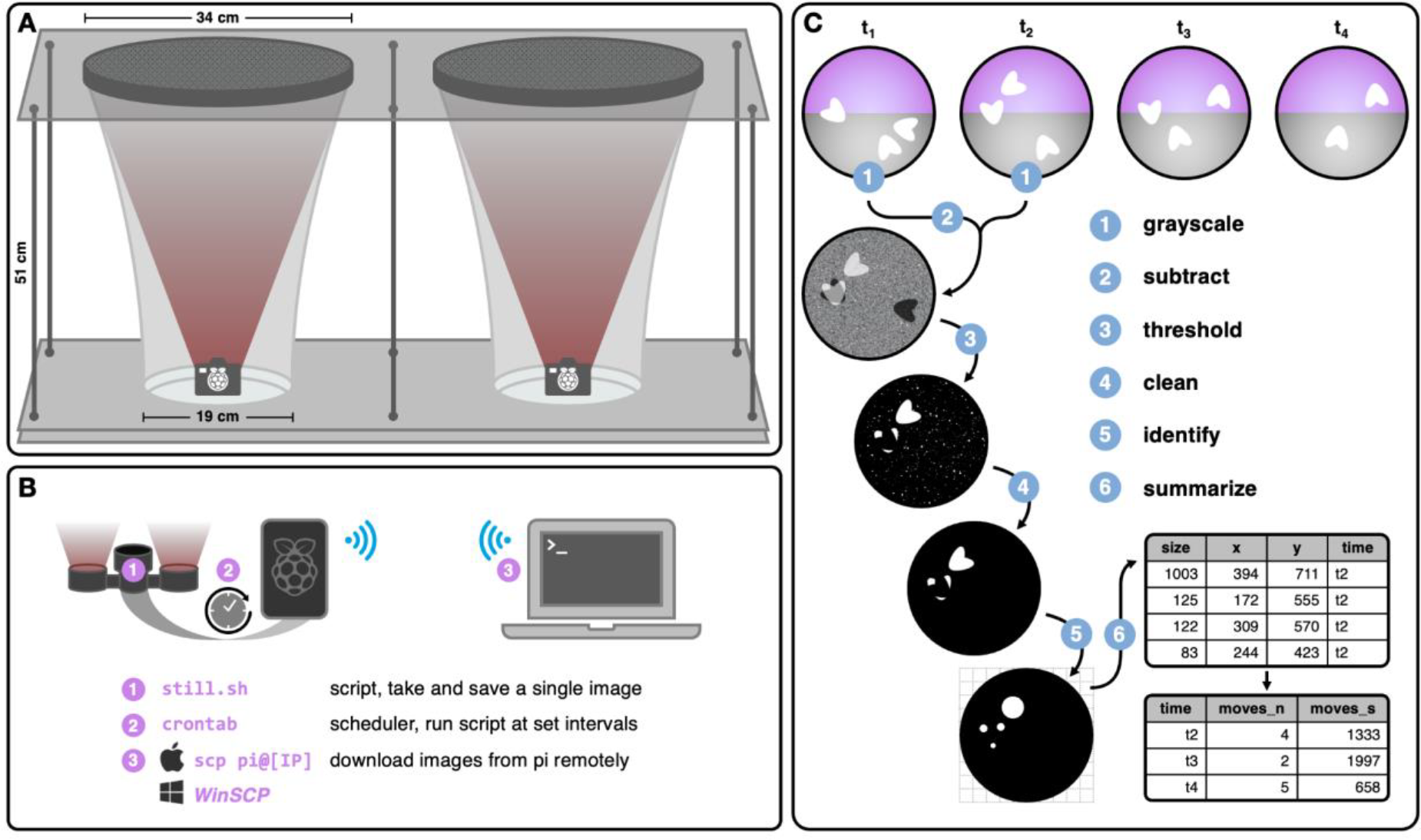
Overview of iLAM design and workflow. (A) Schematic of DIY paired flight cage setup with Raspberry Pi and infrared cameras installed at the base of each. (B) On-board scripts to capture overhead images and remotely transfer them to a personal computer for analysis. (C) Post-process image segmentation workflow to identify and summarize movements between pairs of images.

Although the flight cage design is flexible, we constructed side-by-side pairs of iLAMs from items commonly found at local hardware and art stores ($150 USD/paired setup; Figure 1A). This modular design facilitates disassembly and background customization to increase the detectability of black-(e.g., fireflies) versus white-bodied (e.g., moths) insects under infrared (IR) illumination. The tapered netting also increases detection by limiting movement outside the field-of-view of the camera. At the bottom of each flight cage, one Raspberry Pi Zero W computer and night-vision camera with infrared accessory lights photographs insect movement overhead (Figure 1A-B).

Custom scripts are scheduled with crontab (Reznick, 1993) to capture and save an overhead image (2592×1944 px) as frequently as every five seconds to a mounted micro-SD card. Saved images can be downloaded remotely for post-process segmentation and analysis.

### 2.2 Post-Process Image Segmentation

Briefly, all image segmentation is conducted in R (v.4.1.0, R Core Team) using `imager` (Barthelmé & Tschumperlé, 2019), an R wrapper that integrates the speed of C++ image-processing library `cimg` (Tschumperlé, 2012) with existing R tools. The iLAM wrapper function *find_movements()* grayscales, optionally blurs, and subtracts consecutive images to identify the number, size, location, and timing of all prominent between-image differences (Figure 1C). Only pixel differences greater than a given quantile threshold (99.9%) are retained, as determined by *find_threshold()*; these pixel differences correspond to animals that are absent from the first image (t_i_) and appear in the second image (t_i+1_). Filtered pixels are cleaned and clustered prior to segmentation into blobs, which are visual representations of movement (Figure 1C). The location and size of these identified movements are output into a CSV file for subsequent filtering. This CSV aggregates all movements identified by each iLAM in an experiment. The associated function *parse_movements()* combines data from this CSV with metadata, including trial date, time, light:dark conditions, treatment, and species. To verify accuracy, iLAM identified movements can be visualized over the original images with *plot_movements()* and *make_gif()*. Finally, *make_dam_file()* converts these movement data from a CSV into the standard Drosophila Activity Monitor (DAM) format for activity analyses in `rethomics` (Geissmann, Garcia Rodriguez, Beckwith, & Gilestro, 2019). Users can choose whether to quantify activity as the total number of discrete movements or the summed size of all movements at a given time point.

### 2.3 Case Studies

As a proof-of-concept, we used the iLAM to measure diel activity and circadian phenotypes in three case studies: interspecific variation in the temporal niches of male fireflies (*Photinus greeni, P. marginellus, P. obscurellus*), sex-specific differences in moth diel activity (*O. nubilalis*), and ecotype differences in moth circadian phenotypes (*O. nubilalis*: univoltine, bivoltine). We also compared our measurements of *O. nubilalis* diel activity with those obtained from two TriKinetics LAM25Hs (Waltham, MA), hereafter referred to as DAMs. All experiments were conducted in a climate-controlled room (23.5°C, 40% RH). Following a day of acclimation in the flight cages, insects were recorded for three full days of 16L:8D hours. For circadian phenotyping, *O. nubilalis* moths were additionally exposed to four days of continuous darkness (DD). Further information is provided within the Supplementary Material.

### 2.4 Data analysis

All analyses and visualization utilized the `rethomics` framework (Geissmann et al., 2017) in R (v. 2.4.2, R Core Team 2022). Activity was estimated as the sum, in pixels, of all movements identified between consecutive images taken two minutes apart. These data were combined into 30-minute bins and normalized across individual cages by setting the average activity level for all 30-min bins equal to one. Times were converted to Arbitrary Zeitgeber Time (AZT), where AZT 0 indicates when lights were turned on.

Per Kostadinov et al., 2021, daily activity was smoothed with a Butterworth filter and the *find_peaks()* function from `pracma` (Borchers, 2021) used to identify the timing of peak activity across three days of light:dark (LD) entrainment. We defined nocturnal activity (“nocturnality”) as the percentage of total activity that occurred during scotophase each day (Schlichting & Helfrich-Förster, 2015). Over four days in free-running continuous dark (DD) conditions, endogenous period length and rhythmic strength were estimated using a chi-square periodogram, where period length was the peak value above the significance threshold at α= 0.05 (for more details, see Cai, Hidalgo Sotelo, Jackson, & Chiu, 2022).

## 3. Results

Under 16L:8D, the timing of peak activity was similar across all three firefly species: *P. greeni* (AZT: 17.0 hr, 95% CI: 16.2–17.8), *P. marginellus* (17.0 hr, 95% CI: 17.0–17.0), and *P. obscurellus* (17.2 hr, 95% CI: 16.7–17.6). Similarly, there were no strong differences in nocturnality between *P. greeni* (0.66, *N* = 3) or *P. marginellus* (0.52, *N* = 3). However, there was weak evidence that *P. obscurellus* (0.75, *N* = 3) exhibited greater nocturnality than *P. marginellus* (t = 3.1, df = 2.4, P = 0.07).

In contrast, *O. nubilalis* female (0.95, 95% CI: 91.8–99.0) and male (0.92, 95% CI: 0.84–1.0) moths were almost exclusively nocturnal. Peak diel activity of *O. nubilalis* females (AZT: 19.2 hr) occurred substantially earlier than that of males (22.0 hr; t = 3.95, df = 2, P = 0.05; Figure 2B). This difference persisted when the activity of females (16.8 hr, 95% CI: 16.2–17.4) and males (20.6 hr, 95% CI: 18.1–23.0) was measured with a TriKinetics DAM. However, the timing of females’ peak activity significantly differed between the iLAM and DAM setups (t = 7.41, df = 4.9, P < 0.001).

**Figure 2.**
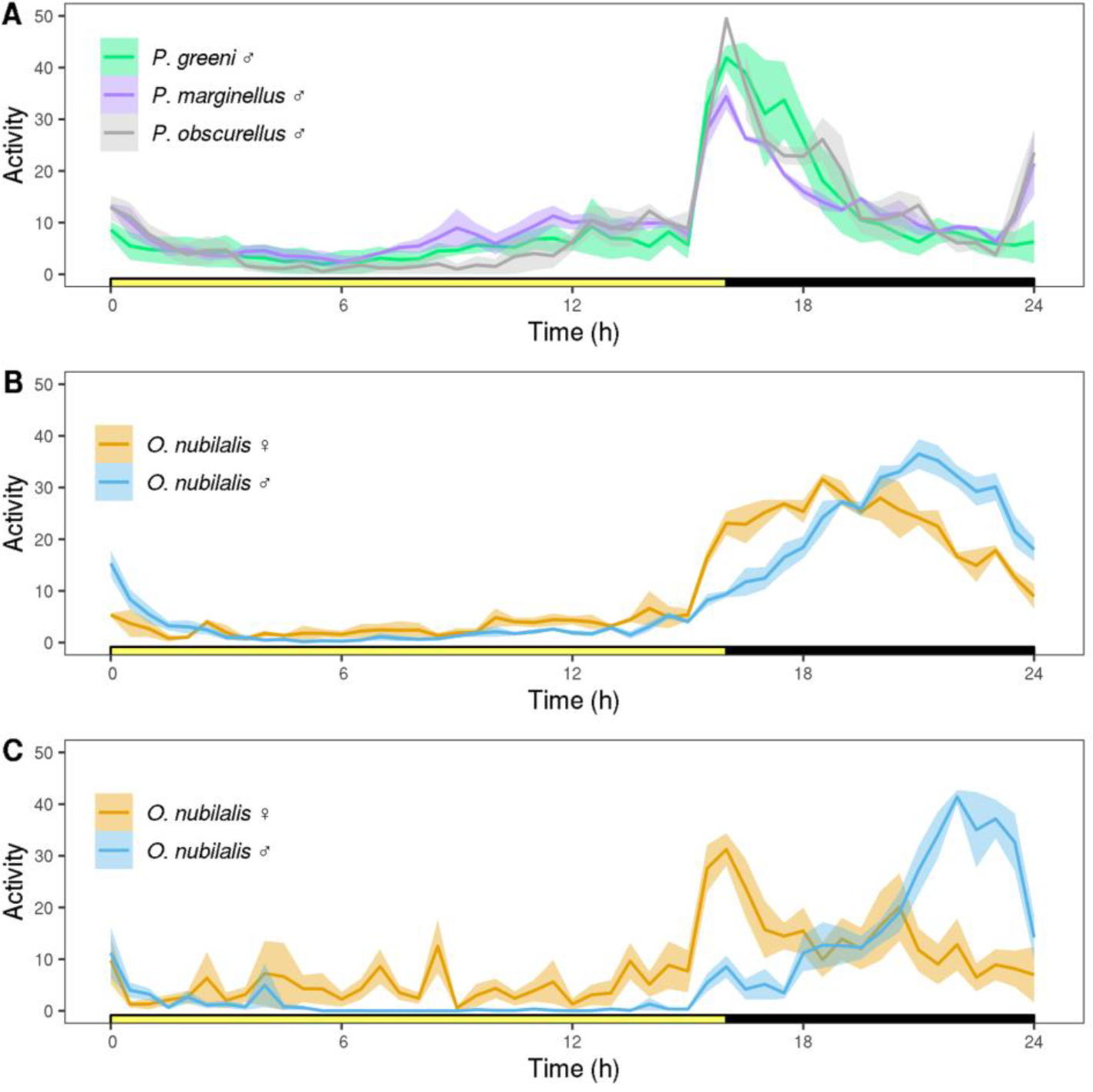
Diel activity of adult *Photinus* fireflies (A) and bivoltine *Ostrinia* moths as measured by iLAMs (B) and TriKinetics DAMs (C). Activity on the y-axis corresponds to the daily average (± SE) of normalized 30-min binned activity. The x-axis shows Arbitrary Zeitgeber Time (AZT); AZT 0 indicates when lights were turned on.

In complete darkness (DD), the rhythmicity of female moths markedly declined (Table 1). In contrast, all male moths remained rhythmic. Univoltine males exhibited a significantly shorter endogenous period length (21.3 hr) than males from a bivoltine genetic background (22.7 hr; t = 6.33, df = 4, P = 0.003; Table 1).

**Table 1.**
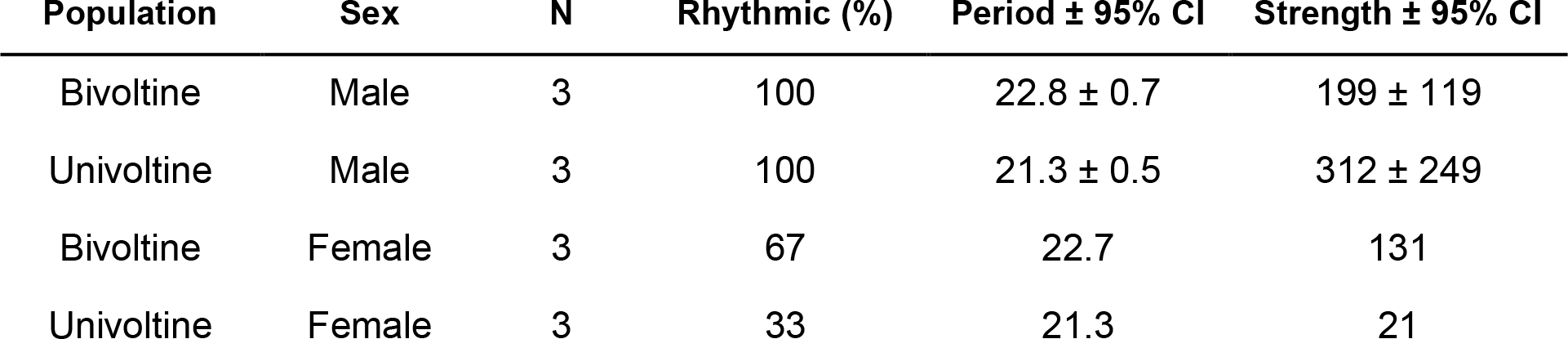
Activity rhythms of *Ostrinia nubilalis* moths from different ecotypes. Activity rhythm in constant darkness is given in hours ± 95% confidence interval.

| Population | Sex | N | Rhythmic (%) | Period ± 95% CI | Strength ± 95% CI |
| --- | --- | --- | --- | --- | --- |
| Bivoltine | Male | 3 | 100 | 22.8 ± 0.7 | 199 ± 119 |
| Univoltine | Male | 3 | 100 | 21.3 ± 0.5 | 312 ± 249 |
| Bivoltine | Female | 3 | 67 | 22.7 | 131 |
| Univoltine | Female | 3 | 33 | 21.3 | 21 |

## 4. Discussion

The iLAM is an affordable and powerful system for monitoring the locomotor activity of large-bodied insects (Figure 1). Even with a small sample size – three paired cages with five to ten individuals each and recording durations of less than a week – our iLAMs proved capable of effectively capturing clear differences in activity timing between species, sexes, and populations (Figure 2, Table 1).

The iLAM produced strikingly similar results to a TriKinetics DAM, the current standard (Figure 2B-2C). The main difference is that the startle effect at the light:dark transition was more pronounced in DAM data (Figure 2C); this may be an artifact of the small diameter of the tubes. DAM data also had higher overall variance because insects were monitored individually instead of in small groups. Regardless, the circadian phenotypes captured here (Table 1) were remarkably consistent with those previously obtained by the same TriKinetics DAMs for bivoltine (period = 22.7 hr) and univoltine (period = 21.4 hr) males as reported by (Kozak et al., 2019).

Unlike existing movement trackers, including DeepLabCut (Mathis et al., 2018), MARGO (Werkhoven, Rohrsen, Qin, Brembs, & de Bivort, 2019), and others (Geissmann et al., 2017; Jolles, 2021), the iLAM is expressly intended to function as an activity monitor. Because it aggregates data from groups of insects and disregards individual movement paths, the iLAM is a highly accessible tool for all researchers with access to a personal computer. Nevertheless, the iLAM can identify the location and size of all movements made by insects within the range of view of the camera. These data are much richer than the current standard: counts of beam interruptions made by insects in a tube (see also Sondhi et al., 2022). Our enclosures, which are large enough to accommodate flight, also permit a broader range of ecologically relevant behaviors. Although not reported here, future studies could easily connect the location and size of identified movements with particular behaviors (Gilbert et al., 2022) such as feeding (Fenske, Nguyen, Horn, Riffell, & Imaizumi, 2018), orientation (Wan, Hayden, Iiams, & Merlin, 2021), calling (Gao et al., 2020), or eclosion (Zhang, Markert, Groves, Hardin, & Merlin, 2017). Because the iLAM saves each image taken during the experiment, this leaves a strong record that can be checked for accuracy and revisited at any point for supplementary analysis.

In the 21st century, natural environments are changing at an unprecedented rate. Now more than ever, there is a critical need to document how organisms alter their activity timing in response to anthropogenic stressors across the seasonal cycle (Gilbert et al., 2022). Accessible, modular tools like the iLAM can catalyze faster adoption of automated activity monitoring, especially for novel study systems that require a more flexible approach. In this way, our simple system has the capacity to dramatically advance the growing field of chronoecology.

## Acknowledgements

We thank Michele and James Dayton for feedback on flight cage design, Vaidehi Chotai and Ross Vieira for feedback on our scripts, Christina Divoll for mounting remote directories, Erik Dopman for providing access to TriKinetics LAMs and corn borer populations, and Sara Lewis for providing space to conduct our experiments. The authors acknowledge the Tufts University High-Performance Computing Cluster (https://it.tufts.edu/high-performance-computing) which was utilized for the research reported in this paper.

## Supplementary Methods

### 2.3.1 Demonstration: Interspecific Differences in Diel activity

In summer 2022, male *Photinus greeni, Photinus marginellus*, and *Photinus obscurellus* fireflies were collected from the wild. *P. obscurellus* (N = 30) were collected from Smith-Andover Field in Lincoln, MA, on June 14th (42.42568, -71.30675; sunrise: 5:08; sunset: 20:22), *P. greeni* (*N* = 30) from Estabrook Woods in Concord, MA, on June 23rd (42.48515, - 71.34635; sunrise: 5:10; sunset: 20:24), and *P. marginellus* (*N* = 46) from Muster Field in Lincoln, MA on August 6th (42.407523, -71.330734; sunrise: 5:43; sunset: 19:57). Species identity was determined from body size, elytral pigmentation, and courtship flash pattern (Lloyd 1969); sex was determined from lantern morphology.

Males were haphazardly divided into groups of approximately five individuals and kept outside until the following day. They were then moved indoors to a climate-controlled room (23.5°C, 40% RH) and evenly distributed among three unpaired flight cages, each supplied with an *ad libitum* source of water and two cups of moist sponges for shelter. To remain consistent with the natural timing of sunrise and set, overhead lights automatically turned on at 05:00 and off at 21:00. Diel activity was recorded over three full days of 16L:8D hr, beginning the day after males were introduced to flight cages.

### 2.3.2 Demonstration: Sex and Population-Specific Differences in Diel activity

Bivoltine European corn borer (ECB) adults were collected from a laboratory population at Tufts University (Medford, MA) that has repeatedly been selected and studied for bivoltine characters (i.e., short post-diapause development timing) and studied previously (Levy et al. 2018 ; Kozak et al. 2019). This population was originally derived from larvae collected in Geneva, NY (42.8680° N, 76.9856° W) and out-crossed with bivoltine ECB from Hollis, NH (42.7668° N, 71.6000° W). Similarly, univoltine ECB adults were sourced from a laboratory stock population, founded from Bouckville, NY (42.8892N, 75.5513), and selected for univoltine characters (Levy et al. 2018 ; Kozak et al. 2019). ECB larvae were reared on artificial corn borer diet (Southland Products, USA) under 16:8 LD at 23.5°C. Pupae from each population/sex (*N* = 15-22) were isolated into individual plastic cups with moist dental wicking. Following eclosion, 1-2 day old adults were randomly selected, sexed, and evenly evenly distributed among three flight cages. All experiments were conducted in a climate-controlled room (23.5°C, 40% RH). Beginning the day after cage introduction, insects were entrained for three full days of 16L:8D hours. *O. nubilalis* were subsequently exposed to four days of continuous darkness (DD) for circadian phenotyping. Each cage contained one *ad libitum* water source and two cups of moist sponges.

### 2.3.3 Demonstration: Comparison with TriKinetics LAM25H

Bivoltine European corn borer adults were singly loaded into 30 TriKinetics LAM25H tubes (25mm diameter) and confined by cotton plug. For a water source, each tube contained a moist cotton ball and hydrated water crystals at the base of each tube. Activity was quantified as the number of beam interruptions that occurred within each 2-minute interval. Recordings occurred in a climate-controlled room (23.5°C, 40% RH) under a 16L:8D cycle. Beginning the day after tube introduction, insects were entrained for three full days of 16L:8D hours. *O. nubilalis* were subsequently exposed to four days of continuous darkness (DD) for circadian phenotyping. However, due to high mortality and low activity, circadian phenotypes could not be quantified from LAM25H insects.

